# Disparities in climate risk-mitigation and recreation services in Grenoble (France)

**DOI:** 10.1101/2025.09.19.677080

**Authors:** Margot Neyret, Samuel Lormel, Alberto Gonzalez-Garcia, Dan Richards, Yves Schaeffer, Mihai Tivadar, Améline Vallet, Sandra Lavorel

## Abstract

Urban green spaces provide cultural and climate risk-mitigation ecosystem services (ES) critical for the well-being of urban residents. Yet these benefits are not equally shared across social groups, and environmental segregation - i.e., the spatial separation of population groups between areas of contrasted environmental quality - is common. For instance, environmental justice studies have shown that green space acreage and quality are typically lower in low-income neighbourhoods. However, more detailed assessments integrating a range of ES and the diverse needs of multiple groups are needed to effectively inform equitable urban planning. Here, we combine socio-demographic spatial data with ES modelling to assess social disparities in the distribution of three ES - heatwave mitigation, flood regulation, and urban recreation - in the Grenoble metropolitan area, France. Our results show that low-income households generally benefit from lower climate risk mitigation, but from equal or slightly greater urban recreation opportunities. We also identify mismatches between ES supply and demand from vulnerable groups, which are particularly acute in disadvantaged neighborhoods. These findings highlight the importance of integrating social needs and vulnerabilities into urban greening strategies. Policies that move beyond generic measures of green space provision and explicitly consider multiple ES can help ensure that urban green infrastructure delivers fair and effective benefits across diverse communities.

## Introduction

The world’s cities currently host 55% of the global population, a share expected to rise to 64% by 2050 (United Nations 2018). This rapid urbanisation comes with multiple environmental management challenges: urban areas have high demand for ecosystem services (ES) due to high population densities, while urban pressure on land often reduces both the supply and accessibility of ES (Lapointe et al., 2020). Urban green areas contribute important ES that support many needs of urban dwellers, but cities also rely on ES from outside their boundaries, such as food production or water from nearby natural and agricultural areas (González-García et al., 2020). Benefits include improved environmental quality (e.g., air pollution abatement) and mitigation of environmental risks such as heatwaves and floods (Andersson et al., 2022), which are expected to become more critical as extreme climate events intensify. A major contribution of urban green spaces to human quality of life is the provision of opportunities for outdoor recreation and relaxation (Meyer-Grandbastien et al., 2020). Time spent in green spaces promotes mental (Marselle et al., 2020) and physical health (Aerts et al., 2020) especially in young people (Chawla, 2015), reduces thermal discomfort during heatwaves (Arnberger et al., 2017) and fosters social relations (Reyes-Riveros et al., 2021). Conversely, limited access to nature and ES among urban dwellers has been linked to growing nature disconnection (Lapointe, 2020; Martín-López et al., 2012), with negative impacts on mental and physical health (Colléony et al., 2020) and overall quality of life (Lapointe et al., 2020).

Cities display wide variation in both environmental conditions and socio-economic structures, often resulting in major environmental disparities. Environmental justice first emerged as a protest against the higher exposure of deprived communities to pollution (Kato-Huerta and Geneletti, 2022; Schlosberg, 2013). As an academic field, it has now expanded its focus to other environmental (dis)amenities (Calderón-Argelich et al., 2021). In urban contexts, disadvantaged groups tend to experience both greater exposure to disamenities and fewer benefits. For instance, people with lower income or minority status often have reduced access to urban green and blue areas (Lapointe et al., 2020). This may result from differences in the availability, quality, or safety of such spaces across neighbourhoods (Rigolon, 2017; Yasumoto et al., 2021). These populations also receive fewer ES related to climate adaptation (Herreros-Cantis and McPhearson, 2021). In cities, such disparities are likely to be related to residential segregation - i.e., the spatial separation of population groups - between areas of contrasted environmental quality - unless explicitly addressed by specific social and planning policies (Law et al., 2022). Understanding how environmental segregation affects multiple dimensions of urban ecosystems’ contributions to well-being, including regulation of environmental hazards and non-material aspects such as recreation, could provide key insights for urban planning and for the development of sustainable and just green infrastructure. Yet, few studies have conceptualised ES supply in urban areas in terms of environmental segregation, particularly when analysing multiple ES (Romero et al., 2012; Schaeffer and Tivadar, 2019; Yi et al., 2019).

Assessing the (un)equal distribution of environmental bads and goods between groups is central to the distributional dimension of environmental justice. However, an equal spatial distribution of ES does not necessarily equate to fair outcomes. For example, ES supply often decreases in densely built areas, which are also highly populated, leading to mismatches between ES supply and demand. Besides, individuals and groups have different ES needs, which shape the environment’s contributions to their well-being. Groups perceive and prioritise different ES (Özgüner, 2011): for instance, Kabisch and Haase (2014) showed that in Berlin green spaces, native German families favoured grassy areas and playgrounds for recreational activities, while older and immigrant families preferred areas for social activities, such as barbeque areas. Such differences influence how social groups benefit from the same green space. Vulnerable groups may also be more dependent on specific ES. For example, children and elderly people are more exposed to heatwave risks and therefore rely more heavily on heat mitigation ES (Reid et al., 2009), but do not necessarily have access to greater ES supply. These vulnerabilities can be particularly severe in deprived populations, who often lack coping strategies, such as air conditioning or the ability to leave the city to enjoy rural green spaces (Martinez-Harms et al., 2018). Evaluating the distribution of ES and the matches or mismatches between their supply and demand across social groups can inform urban planning policies aimed at reducing environmental inequalities.

Here, we assess disparities in ES across social groups in the Grenoble area, a dynamic metropolitan area in southeastern France. We mapped three critical ES - heatwave mitigation, flood regulation by vegetation, and urban recreation - and assessed potential mismatches between ES supply and demand from specific groups, with the following hypotheses: (H1) climate regulation services supply is lower in more densely populated neighbourhoods, which disproportionately affects vulnerable populations in these areas; (H2) mismatches between ES supply and demand are stronger in low-income neighbourhoods and (H3) poor households experience both lower regulating service provision and access to lower-quality urban green spaces. Urban recreation was mapped using a new model developed for the needs of this study, which combines multiple indicators of urban green space attractivity, recreation opportunity, pollution avoidance, and accessibility.

## Methods

### Study area

The Grenoble metropolitan area (Grenoble Alpes Métropole) in southeastern France is an active and dynamic French metropolitan area, with high ecological and social diversity (Vannier et al., 2016, 2019). Grenoble is surrounded by three biodiverse mountain ranges -Vercors, Chartreuse and Belledonne-which benefit from multiple protection measures through two natural parks and several conservation areas. These mountain ranges delineate three valleys (Drac, Isère and Bièvre) that host agricultural activities but are also under pressure from urban sprawl (Vannier et al., 2019, Desgouttes and Gilbert 2014). The valleys face significant climate risks, particularly flood risks from the Isère and Drac rivers, which drain large catchments of the Alps; and heatwaves that strongly affect dense urban areas located at low altitude (ca. 200-300 m a.s.l.) (Foissard et al., 2024). Topographical constraints have led to high population densities in the valleys, reaching up to 100 inhabitants and 50 housing units per hectare in the densest areas of the metropolitan area (95% percentile of 200 m x 200m census squares (Insee, 2019)).

We focus on all urban zones within the commuting perimeter of the Grenoble metropolitan area, i.e. within a 30-minute drive or about 25 km from the city. We define urban areas according to the Global Human Settlement Index as dense or semi-dense clusters, as well as suburban and peri-urban clusters (Schiavina et al., 2023). This encompasses around 70 municipalities and 200 km2, with a total population of about 500,000 inhabitants in 2019 (Fig. S1). Socio-economically, the area has a higher proportion of white-collar and intellectual professions, and fewer workers involved in industry, compared to the national average (Desgouttes and Gilbert, 2014). Although the share of low-income households is lower than in other French metropolitan areas, disparities remain stark, with most low-income households (including many students with no income) concentrated in the south of the area, and much higher average incomes in the north-east (Decorme and Labosse, 2021).

### Ecosystem services models

#### Recreation in urban greenspaces

##### Identification of urban green spaces

We considered all public green spaces within the studied area, as well as in a 500m buffer surrounding it to avoid edge effects. We then selected all vegetated land use types from OpenStreetMap (OpenStreetMap contributors, 2024) that were likely to be publicly accessible, including public parks, forested or grassy areas and wastelands, while excluding private gardens.

##### Development of a recreation opportunity index for urban green spaces

Existing studies of equity in terms of access to urban green space often focus on green space area per person, or distance to the nearest park. Some studies have also described green space quality through user preferences for specific characteristics (Arnberger et al.,2017; Rigolon, 2017), but without accounting for availability. As a result, few studies integrate both green space quality and availability indicators in urban areas (but see Ibes, 2015; Liotta et al., 2020). The recreation opportunity spectrum (ROS) (Byczek et al., 2018; Clark and Stankey, 1979), originally developed for peri-urban and rural areas, captures four dimensions of recreation potential: attractiveness, recreation opportunities, pollution avoidance, and accessibility. We identified published indicators for each of these dimensions, adapted to urban areas (Fig. 1). Many indicators found in the literature referred to the presence of specific features (e.g. water bodies), without mentioning a distance up to which they enhance surrounding green space quality. In such cases, we applied distance decay functions, with a half-life of 300m which roughly corresponds to a 5 min walking distance (a quarter mile in Rigolon, 2017).

**Figure 1.**
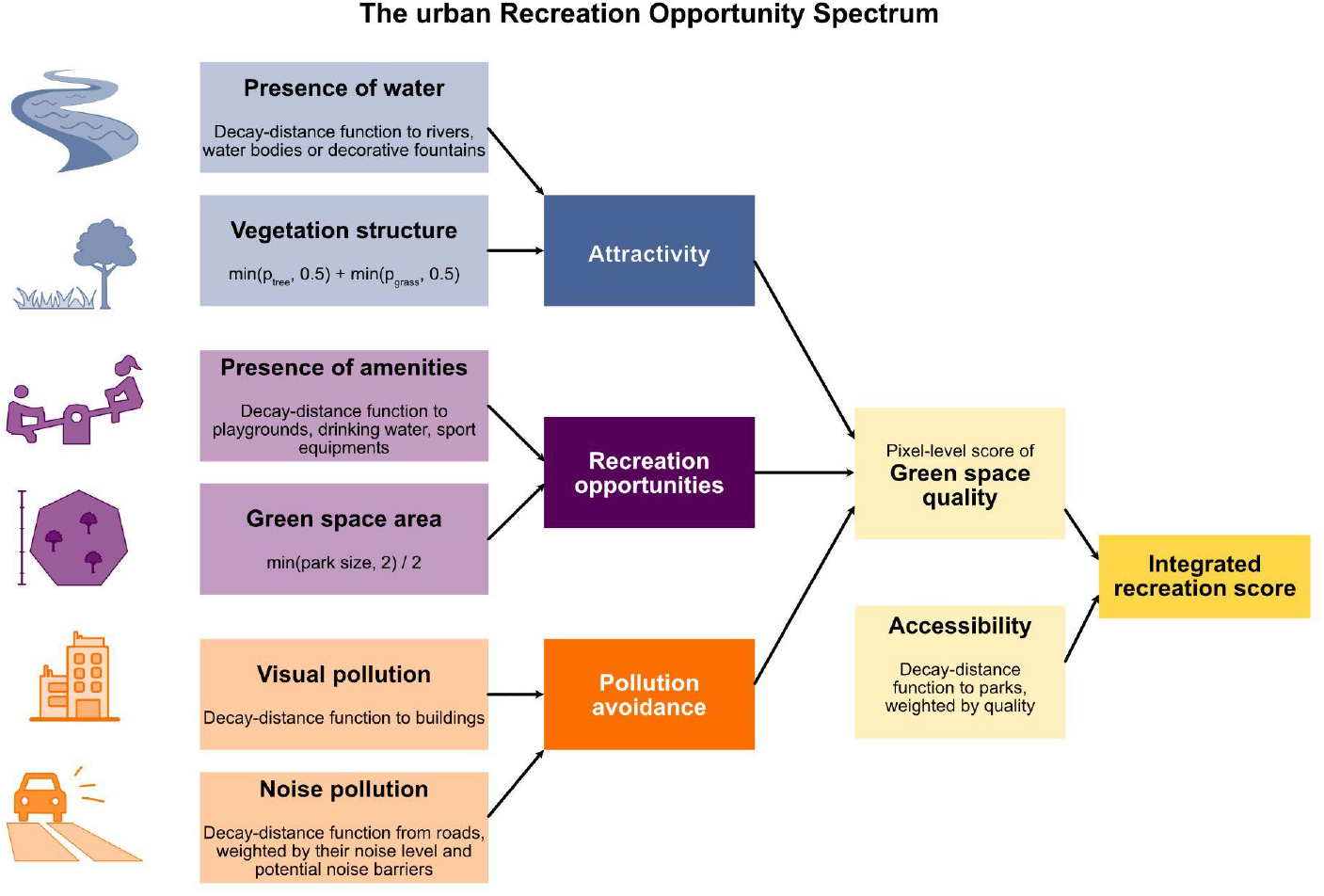
Components of the urban Recreation Opportunity Spectrum developed in this study.

We defined green space attractiveness using two indicators commonly applied in the literature: vegetation diversity and the presence of water bodies. Both indicators were derived from fine-scale land use cover maps provided by the French National Institute of Geographic and Forest Information (IGN, 2023), aggregated at 10 metres resolution. Previous studies have shown that visual appreciation of the vegetation is maximised by a mix of grass and tree cover (Jiang et al., 2015), although they did not specify optimal proportions. We therefore assumed equal proportions to be optimal. The vegetation attractiveness indicator for each green space pixel was measured as:

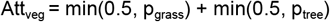

Where p_grass_ and p_tree_ are the proportion of grassy and woody vegetation located in urban green spaces, within a 300m radius from the target pixel. Att_veg_ ranges between 0 and 1, is maximised when the surrounding area contains exactly 50% grass and 50% tree cover, and decreases when non-vegetated areas are present or when the ratio of grass:tree departs from 1:1. The water attractiveness indicator (Attwat) was calculated for each pixel as a distance-decay function with a 300-m half-life from water bodies (rivers, ponds, fountains):

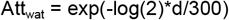

With d the distance (m) from the target pixel to the closest water body. The overall attractivity score was then:

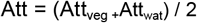

Recreation opportunities were measured from green space size and available recreation infrastructure. Park size strongly influences use and enjoyment (Dade et al., 2020; Sarı and Bayraktar, 2023). Benefits increase up to a size threshold. While 1 ha is often used, our study also included much larger peri-urban green spaces. We therefore chose 2 ha as a threshold, corresponding to the 80th percentile of park size in the study area and consistent with the 1.5–2 ha minimum often cited for adult physical activity (Liotta et al., 2020). Because OpenStreetMap sometimes split green spaces with small roads, we merged spaces within 20 m of each other, assuming they function as one unit. The green space size indicator was calculated as:

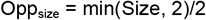

Where size is the area (ha) of the green space. This measure was applied to all pixels within each green space. We also considered three amenities valued by users: playgrounds, sport equipment, and drinking water fountains (Özgüner, 2011), which we extracted from OpenStreetMap. For each amenity type, we calculated pixel-specific scores representing the presence of available amenities, down-weighted by distance, as a decay function with a 300m half-life, and averaged the three scores. The total score of amenities was thus:

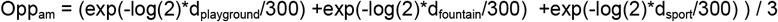

With d_x_ the distance (m) to the closest corresponding amenity. The overall recreation opportunity score was:

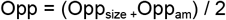

Noise and visual pollution such as building visibility reduce the recreational quality of green spaces (Carvalho and Cleto, 2012). Air pollution may also hinder use; but vegetation’s capacity to mitigate is debated (Xing et al., 2023) and models remain uncertain, therefore we excluded it. Visual pollution was measured as a distance-decay function from the nearest building:

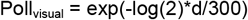

where d is the distance (m) to the closest building. Noise pollution in cities comes mainly from traffic (Carvalho and Cleto, 2012), We estimated it using distance to roads and presence of noise barriers, rather than detailed noise reduction modelling. Roads were assigned noise levels: pedestrian streets and small alleys (65dB), larger city streets (80dB), and highways and railways (90db) (Margaritis et al., 2018). We then applied a cost-distance approach to represent noise barriers: pixels occupied by buildings, which act as strong barriers against noise, had a cost of 50; pixels occupied by woody vegetation had a cost of 10; and pixels with neither had a cost of 1. The noise pollution for each pixel was calculated as

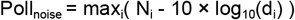

With N_i_ and d_i_ the noise level and distance to the closest road of type i.

The combined pollution score was thus:

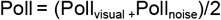

And the overall quality score for each green space pixel as the average of the attractiveness, opportunity, and inverted pollution scores:

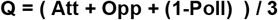

##### Access to urban green spaces

The last dimension of the Recreation Opportunity Spectrum is green space accessibility. We accounted for the fact that nearby, average-quality spaces used on a daily basis may provide similar value as larger, distant, higher-quality ones. For this we applied a distance weighting to green space quality. For each pixel, we calculated the average quality scores in a 1 kilometre buffer, down-weighted by distance using a 300-m half-life decay function.

#### Heatwave mitigation

Heatwave mitigation (HMI) was calculated using the inVEST software (Natural Capital Project, 2024) which assesses the contribution of vegetation to decreasing urban temperatures during heat waves. The model considers evapotranspiration, albedo, shade coefficients for each land cover type as well as distance to vegetation. It was run using the EUSALP land cover map (Marsoner et al., 2023), CHELSA climate (Karger et al., 2017) and evapotranspiration data (Zomer et al., 2022). Parameters were adapted from nearby case studies (Bosch et al., 2021); details on data sources can be found in the supplementary methods.

#### Flood regulation

Flood regulation was measured using inVEST Urban Flood risk mitigation model, which assesses the potential for excess water retention based on soil hydrologic group, land cover and topography. The model applies the ESDA curve number approach to calculate the water saturation level for each land cover type within each soil group. It was run using EUSALP land cover map (Marsoner et al., 2023), CHELSA climate (Karger et al., 2017) and hydrological soil typology data (Ross et al., 2018). Details on data sources can be found in the supplementary methods.

### Socio-demographic data

Socio-demographic variables were obtained from the open Filosofi dataset from the French National Institute of Statistics and Economic Studies (Insee, 2019). This dataset provides information on age structure and economic status at a 200-m grid, based on fiscal and social administrative sources from 2019.

We focused on population density, proportion of poor households (income per consumption unit. <60% of the national median; hereafter *poor households*; others are *secure households*), and living standards as indicators of economic status. The proportion of children (<14) and elderly (>65) were indicators of ES demand. In the study area, poor households represented 13% of all households, while elderly and children represented 20% and 13% of the total population, respectively.

### Data analysis

All analyses were conducted in R (v. 4.4.1, R Core Team, 2024). Key packages included sjstats, spatialreg, terra, boot, OasisR (Angelo Canty et al., 2024; Bivand et al., 2024; Hijmans et al., 2024; Lüdecke, 2024; Tivadar, 2019).

We analysed ES disparities in relation to socio-economic status using two complementary variables: (1) the number of poor versus secure households, and (2) the average living standard, calculated as the sum of living standards of all individuals in each 200 m square divided by the number of individuals.

First, we compared the mean ES supply between poor and secure households, weighting each 200m square supply by the number of households in each category. 95% confidence intervals were calculated with a bootstrap approach with 999 iterations. Alternatively, we calculated a Gini-based environmental segregation index (GINIes), inspired from environmental centralisation indices (Schaeffer and Tivadar 2019). GINIes measures the degree of unevenness of the relative spatial distribution of two groups x and y, compared to the distribution of an amenity score - for instance ES supply.

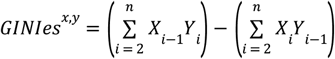

Where, *n* is the number of spatial units and X and Y are the cumulative percentage of each group population through the *i*th spatial unit, spatial units being ordered decreasing by amenity score. Negative values indicate that the first group (x) is more segregated from the amenity than the second group (y).

Schaeffer and Tivadar (2019) proposed a method to test the statistical significance of such indices: the observed value can be compared to an expected value under the null hypothesis that all groups come from the same population. This is done through Monte Carlo simulations (999 iterations), where households are randomly reassigned residential locations, with a location’s assignment probability proportional to its current residential capacity. Pseudo p-values are then calculated as the proportion of simulations with values above (or below) the observed value. This approach provides a robust, distribution-free test of environmental inequality.

To assess whether low-income areas benefit from cumulatively lower ES supply, we discretized each ES into low, medium and high supply, based on 33% quantiles (called low, medium or high supply levels). We then compared proportions of individuals living in areas with at least two ES provided at low level using a two-sided z-test. As complementary analyses, we also ran spatial regression (function errorsarlm) with i) ES supply as response variable, and the proportion of poor households as predictor, weighted by the total number of households (see supplementary material) or ii) the average living standards as response variable and number of services provided at low level as predictor. Spatial autocorrelation in model residuals was considered only for neighbours within 250 m (ensuring inclusion of diagonal pixels) and was significant in all models.

We applied a similar approach to assess potential supply–demand mismatches across the study area. For each ES, we identified socio-demographic variables representing demand. Population density was used as a generic demand indicator for all ES, as it reflects the number of individuals potentially benefitting from the ES in each square. Additional demand indicators were defined for specific ES:

- For heat mitigation, the densities of elderly people and children were used, as both groups are particularly vulnerable to high temperatures.
- For recreation, the density of children was used, as they are major users of green spaces.

As with ES supply, demand variables (the number of individuals, elderly people and children in each square) was divided into 33% quantiles (low, medium, high demand). Each square was classified by its joint demand and supply levels. We further compared mismatches across socio-economic contexts by splitting the study area into high- and low-living-standard squares (above or below the median). We then compared the proportion of individuals benefiting from low ES supply across these areas using two-sided z-tests.

## Results

### 1. Spatial variability in ecosystem services

Heat mitigation and flood mitigation were particularly low in the most densely populated areas of the region, which correspond to the built-up city centre (Fig. 2a-c). The distribution of urban recreation opportunities was more complex: while vegetation cover was much higher in the less densely populated fringes (see Fig. S2), the quality and density of public urban green spaces was quite high across the entire area, including in densely built neighbourhoods (Fig. 2d).

**Figure 2.**
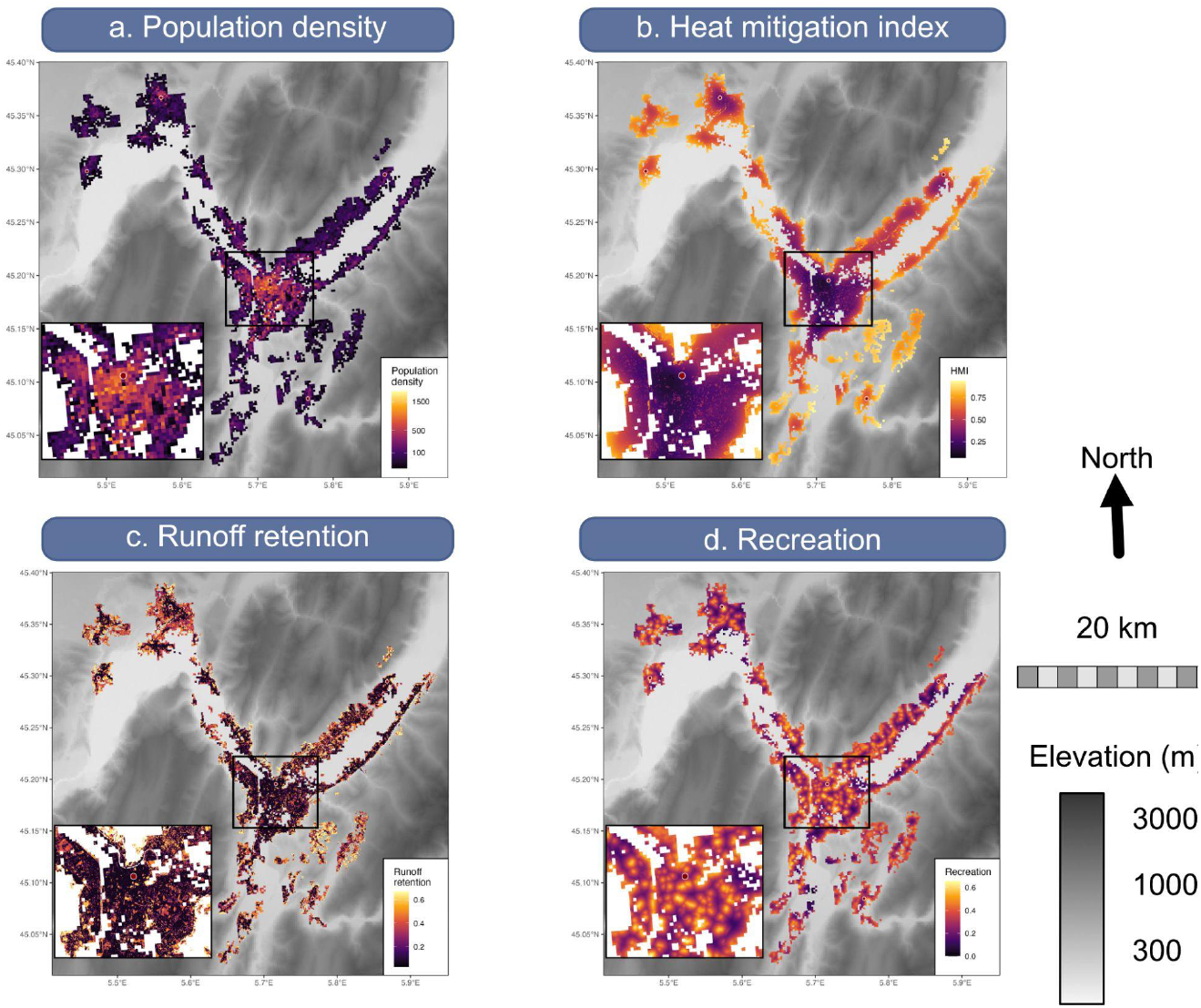
Variation in population density (a), heat mitigation (b), flood mitigation (c) and recreation in urban green spaces (d) across the study area. The inset shows the densest area of the metropolitan area.

### 2. Economic disparities in ES benefits

Overall, poor households and areas with lower living standards experienced lower levels of heat mitigation and flood mitigation, but had relatively good access to urban recreation services.

On average, poor households had lower heat mitigation (0.357 [95% confidence interval: 0.356–0.366]) compared to secure households (0.422 [0.413–0.433]). Flood mitigation showed a similar, though weaker pattern (poor: 0.111 [0.105–0.117], secure: 0.136 [0.131–0.143]) (Fig. 3a–b). These results reflect the concentration of poor households in densely populated areas with limited ES provision (Fig. 2). Conversely, poor households benefited from slightly higher recreation opportunities, although the difference was minimal and confidence intervals overlapped (poor: 0.33 [0.327–0.343], secure: 0.32 [0.130–0.329]; Fig. 3c). Spatial regression analyses mostly supported these results (see Supplementary Results).

**Figure 3.**
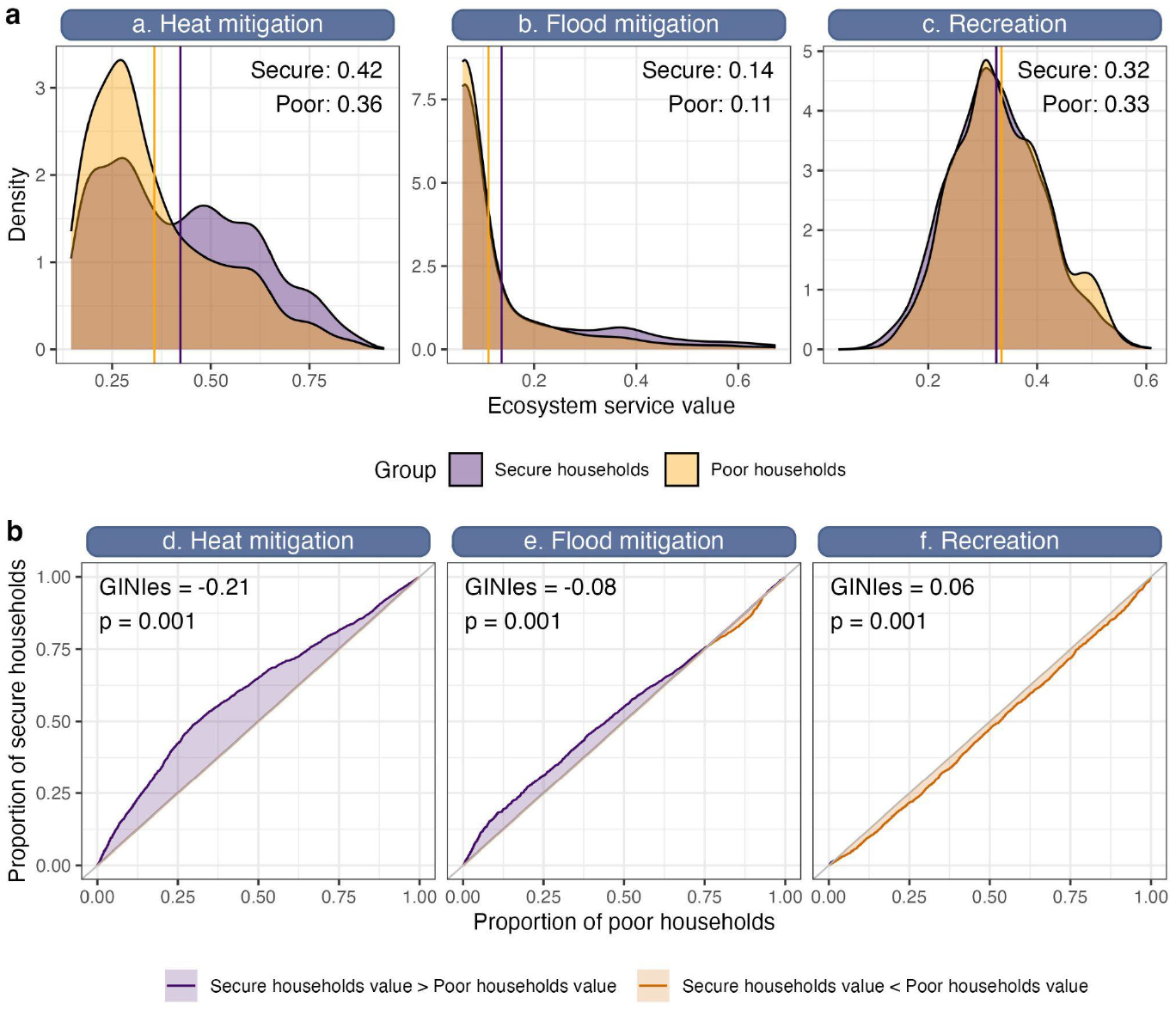
**(a-c)** Comparison of the distribution of ES supply for poor and secure households. Values correspond to the average ES supply, weighted by the number of poor or secure households per square. **(d-e)** Environmental segregation curves showing the cumulative proportion of poor (x-axis) and secure (y-axis) households when spatial units are ordered from highest to lowest ES score. The Gini based Environmental Scores Inequality Index measures the degree of unevenness of the relative spatial distribution of poor and secure households compared to the distribution of each ES. Higher values (purple line above the diagonal), indicate that secure households are less segregated from the ES than poor households, and vice versa.

Environmental segregation analyses confirmed these findings. Poor households had stronger environmental segregation than secure households for heat mitigation (GINIes = −0.21, pseudo p-value ≤ 0.001) and to a lower degree for flood mitigation (GINIes = −0.08, pseudo p-value ≤ 0.001). Conversely, secure households were slightly more segregated from recreational opportunities than poor households (GINIes=0.06, pseudo p-value ≤ 0.001).

Most residents thus had limited access to environmental amenities, but poor households also tended to accumulate multiple deficits. Significantly fewer secure (54%) than poor households (63%, P < 0.001) live in areas where at least two ES are provided at a low level. Consistently, when accounting for spatial autocorrelation, the living standard of people living in areas with at least two ES provided at low level was significantly lower (23.8 ± 4.4 k€) than in other areas (25.8 ± 1.6 k€, P < 0.001).

### 3. Supply-demand mismatches

The low supply of regulating services in densely populated areas (Fig. 2) led to pronounced supply–demand mismatches (Fig. 5a). Overall, 67% of the population lived in areas with low heat mitigation, and 61% in areas with low flood mitigation. Focusing on vulnerable groups, children were equally segregated from the heat mitigation service as the rest of the population (GINIes = 0.003, insignificant pseudo p-value = 0.15). Conversely, elderly people were slightly less segregated from heat mitigation than the rest population (GINIes = 0.03, pseudo p-value = 0.017), which could result from active relocation of older and wealthier people to cooler areas, or historical legacies of longer-term residents living in more vegetated areas which have not been transformed by recent projects. Regarding recreation, 32 % of children lived in areas with low recreation (Figure 5a); and despite higher needs they were more segregated from this amenity than the rest of the population (GINIes = 0.013, pseudo p-value ≤ 0.001).

**Figure 5.**
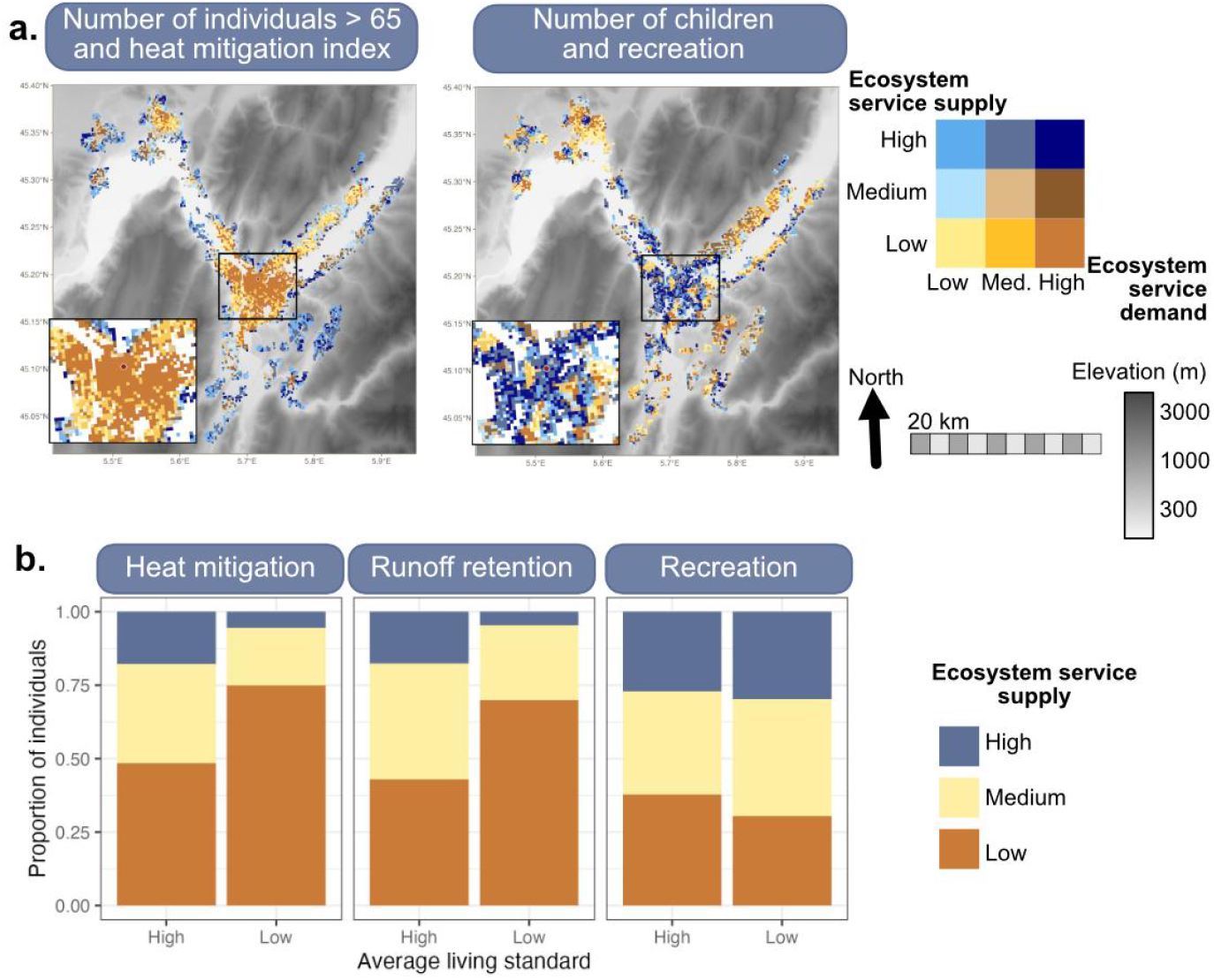
ES supply-demand matches and mismatches. a. Mismatch between demand and supply of heat mitigation: most areas with high density of people above 65 years old have low supply of heat mitigation (right; orange areas); match between demand and supply recreation: many areas with high children density benefit from high recreation supply (dark blue). B. Mismatches are higher in areas with low living standards for regulating services but not recreation. Plots show, in proportion, the number of individuals (as an indicator of demand) living in areas with high (respectively low) average living standards benefiting from low, medium or high service supply.

The magnitude of these mismatches changed with living standards (Fig. 5b). For heat mitigation, the mismatch was stronger in areas with low average living standards: 74% of the overall population (74% of people above 65 years and of children) living in areas with low average living standards benefit from very low heat mitigation, while this was the case only for 48% of the overall population (resp. 51% and 45% for people above 65 and children) in areas with high living standards (all p ≤ 0.001). Conversely, the mismatch for recreation was slightly lower in areas with low living standards, as 29% of people (31% of children) in areas with low living standards benefit from a high recreation service, against 27% in areas with high living standards.

## Discussion

### Contrasting patterns for regulating and recreation services

In our study area, climate risk regulation services were particularly low in densely populated areas in the core of the Grenoble metropolitan area, where vegetation cover is limited. These areas also had lower average living standards and a higher proportion of poor households, leading to strong socio-economic disparities in access to regulating ES. Mismatches between ES supply and demand were especially pronounced in low-income areas. This is particularly problematic, as residents of these areas may accumulate multiple vulnerabilities (Aznarez et al., 2024) and have reduced capacity to cope with or adapt to challenging environmental conditions. For instance, they may face poorly insulated housing or have fewer transport options to visit cooler areas or leave the city during heatwaves (Gronlund, 2014). Engineered solutions can provide alternative approaches to mitigate climate risks; for instance, flood risk can be mitigated by grey infrastructure such as sewers or dykes. We did not assess the distribution of these engineered mitigation options across social groups, but previous work shows that green infrastructure can significantly and cost-effectively reduce environmental risks (Chen et al., 2021; González-García et al., 2025), and should therefore be considered in future efforts to mitigate current inequalities. Our findings nevertheless highlight the likely inadequacy of current green infrastructure in deprived areas of our study region.

Low-income urban residents are often those most affected by nature disconnection and lack of access to nature (Richards et al., 2023). With little access to private gardens or natural areas outside the city, these populations rely heavily on public parks (Tu et al., 2016). We found that recreation services from public urban green spaces tended to be equal or higher in denser, more deprived areas compared with wealthier neighbourhoods. This contrasts with earlier studies. In Paris, green spaces in poorer suburbs are often degraded wastelands with few amenities (Liotta et al., 2020). Similar patterns of reduced availability, attractiveness, and accessibility of urban green spaces for disadvantaged groups have been reported in Germany (Kabisch and Haase, 2014), Austria (Neier, 2023), and many Central and Eastern European cities (Kronenberg et al., 2020). Two factors might explain this discrepancy. First, our approach integrated not just the amount of green space, but also their quality, amenities, and accessibility. This suggests that dense, lower-income areas benefited from well-equipped public parks in relatively high numbers. Such patterns may reflect historical and more recent urban planning policies in Grenoble, including the creation of “eco-neighbourhoods” with high-quality green spaces in dense urban districts. However, the extent to which these policies reduce social disparities remains debated, especially as they often do not integrate targeted actions to tackle environmental or social disparities (Lejemtel et al., 2025). Second, our analysis measured access only in terms of distance. Other critical aspects, such as safety or overcrowding, also condition the recreational value of urban green spaces (Rigolon, 2017; Yasumoto et al., 2021). Finally, our results should be interpreted within the broader context of multidimensional access to nature both within and outside urban areas. Our analysis included only public green spaces in and near the city. In our study region, Schaeffer and Tivadar (2019) showed that poorer households had less vegetation altogether in their surroundings. Yet, in the present study private gardens, which provide major opportunities for outdoor activities and relaxation, but are more common in high-income areas (Venter et al., 2020), were excluded. Wealthier households also likely enjoy easier access to high-quality green areas outside the city (Martinez-Harms et al., 2018), such as nearby rural or mountain landscapes - which are abundant and of high quality in the study region (Byczek et al., 2018), facilitated by their residential location on the metropolitan periphery and/or easier transports such as car ownership. In summary, residents of the poorer neighbourhood in the area thus benefit from abundant and high-quality urban public green spaces, but are likely to face a number of barriers to access other types of green areas we did not consider.

### Considerations for urban planning and policy

Acknowledging the intertwined issues of urban green infrastructure availability, quality and accessibility for contrasted publics raises a significant challenge: how can urban planning provide greater ES to disadvantaged and vulnerable populations, while addressing environmental segregation?

One approach is to improve ES supply by creating and maintaining high-quality, multifunctional urban green spaces that provide a range of benefits such as recreation and environmental risks mitigation. This might be challenging in dense urban areas where space is limited, often requiring major restructuring of the urban fabric. Other nature-based solutions such as green roofs or street trees might be easier to implement but likely provide different sets of ES (Evans et al., 2022). In both cases, what kind of interventions are implemented and where so that the resulting benefits are fairly distributed needs to be considered. Careful, case-by-case evaluation is therefore needed to assess which forms of green infrastructure deliver which benefits, and to whom, to tackle environmental disparities. These interventions should also rely on ecosystems and species resilient to climate change, to ensure long-term ecological and social sustainability (Esperon-Rodriguez et al., 2022). Additional levers to address unequal ES distribution also exist on the demand side for example when deciding where to accommodate disadvantaged households, so they can benefit from green areas and avoid exclusion from newly greened, gentrified neighborhoods (Anguelovski et al., 2022). In Grenoble, for example, some policies reserve housing for low-income households in newly renovated neighbourhoods, which often benefit from quality green areas. To support effective planning, future work should identify priority hotspots for action: areas with low ES supply and high concentration of vulnerable people, or areas with high ES supply where vulnerable people are under-represented. Existing inequity hotspot-identification approaches (Heckert and Rosan, 2016; Schaeffer and Tivadar, 2019) should be extended to include multiple dimensions of well-being (beyond economic factors; e.g., Liotta et al., 2020) and multifunctional outcomes rather than single ES. This should be complemented by participatory processes to co-design just and efficient interventions that support vulnerable groups’ access to nature and environmental equity.

While public actions need to consider both the supply and demand side of environmental disparities, neighbourhoods will continue to be transformed by residential mobilities after the public interventions, and so will environmental inequalities (Nygaard, 2024). A major research priority is therefore to understand the dynamic processes behind these inequalities, which might include both historical patterns either related (e.g. pollution, Heblich et al., 2021) or unrelated to environmental factors. This involves identifying why vulnerable or wealthy people live where they live and, more generally, the mechanisms by which households are spatially sorted in the presence of multiple, unevenly distributed ES. Residential location choices are influenced not only by property prices, but also by proximity to jobs, public transports, facilities and services, social interactions (Gaigné et al., 2022; Goffette-Nagot and Schaeffer, 2013) and preference for specific ES (Pan et al., 2021). Natural amenities in particular affect residential choice, with impacts differing across socio-professional groups (Schaeffer et al., 2016). Addressing the motivations behind residential location choice can thus be another key lever to counter environmental segregation. For instance, current urban planning in the Grenoble metropolitan area aims to improve the quality of life and attractivity of the urban areas to limit urban sprawl (Vannier et al., 2019a) and the outmigration of wealthier households to greener peri-urban or rural settings — an option largely unavailable to low-income households.

## Conclusion

Our study highlighted complex environmental disparities patterns in the Grenoble metropolitan area, with contrasted patterns of disparities and supply-demand mismatches depending on the service considered. In the context of dense European cities and rapidly evolving climate risks, urban greening policies need to explicitly target social disparities - including diversity in access capabilities, vulnerabilities, and preferences - to address environmental segregation and promote just environmental benefits across social groups.

## Acknowledgements

The European Union’s Horizon 2020 research and innovation programme under the Marie Skłodowska-Curie grant agreement No 101104374 provided funding to MN, SLa, SLo.

Project RECONNECT, Biodiversa+, the European Biodiversity Partnership under the 2021–2022 BiodivProtect joint call for research proposals, provided funding to SLa and AG by the ANR Agence Nationale de la Recherche (ANR-22-EBIP-0009-06). DR was supported by Strategic Science Investment Funding for Crown Research Institutes from the New Zealand Ministry of Business, Innovation and Employment’s Science and Innovation Group.

## Authors contributions

MN, SLo, SLa designed the study. AG, SLo, MN provided ES models. YS, MT provided environmental segregation analyses. SLo, MN, YS, MT conducted data analyses.

All authors contributed critically to manuscript writing.

## Data and code availability

The code will be made available on the corresponding author’s GitHub page before publication.

## References

Tivadar M. (2019). OasisR: An R Package to Bring Some Order to the World of Segregation Measurement. Journal of Statistical Software, 89(7), 1–39. doi:10.18637/jss.v089.i07.

Aerts, R., Nemery, B., Bauwelinck, M., Trabelsi, S., Deboosere, P., Van Nieuwenhuyse, A., Nawrot, T.S., Casas, L., 2020. Residential green space, air pollution, socioeconomic deprivation and cardiovascular medication sales in Belgium: A nationwide ecological study. Science of The Total Environment 712, 136426. 10.1016/j.scitotenv.2019.136426

Andersson, E., Haase, D., Kronenberg, J., Langemeyer, J., Mascarenhas, A., Wolff, M., Elmqvist, T., 2022. Based on nature, enabled by social-ecological-technological context: deriving benefit from urban green and blue infrastructure. E&S 27, art18. 10.5751/ES-13580-270418

Angelo Canty, Brian Ripley, Alessandra R. Brazzale, 2024. boot: Bootstrap Functions (Originally by Angelo Canty for S).

Anguelovski, I., Connolly, J.J.T., Cole, H., Garcia-Lamarca, M., Triguero-Mas, M., Baró, F., Martin, N., Conesa, D., Shokry, G., del Pulgar, C.P., Ramos, L.A., Matheney, A., Gallez, E., Oscilowicz, E., Máñez, J.L., Sarzo, B., Beltrán, M.A., Minaya, J.M., 2022. Green gentrification in European and North American cities. Nat Commun 13, 3816. 10.1038/s41467-022-31572-1

Arnberger, A., Allex, B., Eder, R., Ebenberger, M., Wanka, A., Kolland, F., Wallner, P., Hutter, H.-P., 2017. Elderly resident’s uses of and preferences for urban green spaces during heat periods. Urban Forestry & Urban Greening 21, 102–115. 10.1016/j.ufug.2016.11.012

Aznarez, C., Kumar, S., Marquez-Torres, A., Pascual, U., Baró, F., 2024. Ecosystem service mismatches evidence inequalities in urban heat vulnerability. Science of The Total Environment 922, 171215. 10.1016/j.scitotenv.2024.171215

Bivand, R., Piras, G., Anselin, L., Bernat, A., Blankmeyer, E., Chun, Y., Gómez-Rubio, V., Griffith, D., Gubri, M., Halbersma, R., LeSage, J., Li, A., Li, H., Ma, J., Mallik, A., Millo, G., Pace, K., Parry, J., Peres-Neto, P., Rüttenauer, T., Sarrias, M., Sayago, J., Tiefelsdorf, M., 2024. spatialreg: Spatial Regression Analysis.

Bosch, M., Locatelli, M., Hamel, P., Remme, R.P., Chenal, J., Joost, S., 2021. A spatially explicit approach to simulate urban heat mitigation with inVEST (v3.8.0). Geoscientific Model Development 14, 3521–3537. 10.5194/gmd-14-3521-2021

Byczek, C., Longaretti, P.-Y., Renaud, J., Lavorel, S., 2018. Benefits of crowd-sourced GPS information for modelling the recreation ecosystem service. PLOS ONE 13, e0202645. 10.1371/journal.pone.0202645

Calderón-Argelich, A., Benetti, S., Anguelovski, I., Connolly, J.J.T., Langemeyer, J., Baró, F., 2021. Tracing and building up environmental justice considerations in the urban ecosystem service literature: A systematic review. Landscape and Urban Planning 214, 104130. 10.1016/j.landurbplan.2021.104130

Carvalho, A.P., Cleto, R.A., 2012. Sound and noise in urban parks. Proc. Mtgs. Acoust. 18, 040001. 10.1121/1.4772735

Chawla, L., 2015. Benefits of Nature Contact for Children. Journal of Planning Literature 30, 433–452. 10.1177/0885412215595441

Chen, W., Wang, W., Huang, G., Wang, Z., Lai, C., Yang, Z., 2021. The capacity of grey infrastructure in urban flood management: A comprehensive analysis of grey infrastructure and the green-grey approach. International Journal of Disaster Risk Reduction 54, 102045. 10.1016/j.ijdrr.2021.102045

Clark, R.N., Stankey, G.H., 1979. The recreation opportunity spectrum: a framework for planning, management, and research. Gen. Tech. Rep. PNW-GTR-098. Portland, OR: U.S. Department of Agriculture, Forest Service, Pacific Northwest Research Station. 32 p 098.

Colléony, A., Cohen-Seffer, R., Shwartz, A., 2020. Unpacking the causes and consequences of the extinction of experience. Biological Conservation 251, 108788. 10.1016/j.biocon.2020.108788

Dade, M.C., Mitchell, M.G.E., Brown, G., Rhodes, J.R., 2020. The effects of urban greenspace characteristics and socio-demographics vary among cultural ecosystem services. Urban Forestry & Urban Greening 49, 126641. 10.1016/j.ufug.2020.126641

Decorme, H., Labosse, A., 2021. De forts écarts de revenus entre communes au sein des métropoles - Analyses Auvergne-Rhône-Alpes. Insee Analyses Auvergne-Rhône-Alpes.

Desgouttes, S., Gilbert, A., 2014. Grenoble-Alpes Métropole : une agglomération jeune, spécialisée dans les activités scientifiques. Insee Analyses Rhône-Alpes 6.

Esperon-Rodriguez, M., Tjoelker, M.G., Lenoir, J., Baumgartner, J.B., Beaumont, L.J., Nipperess, D.A., Power, S.A., Richard, B., Rymer, P.D., Gallagher, R.V., 2022. Climate change increases global risk to urban forests. Nat. Clim. Chang. 12, 950–955. 10.1038/s41558-022-01465-8

Evans, D.L., Falagán, N., Hardman, C.A., Kourmpetli, S., Liu, L., Mead, B.R., Davies, J.A.C., 2022. Ecosystem service delivery by urban agriculture and green infrastructure – a systematic review. Ecosystem Services 54, 101405. 10.1016/j.ecoser.2022.101405

Foissard, X., Rome, S., Bigot, S., Rousset, E., Fouvet, A.-C., 2024. A new high spatial density temperature dataset in the Grenoble alpine valley (France) for urban heat island investigation and climate services dedicated to municipalities purposes. Data in Brief 55, 110553. 10.1016/j.dib.2024.110553

Gaigné, C., Koster, H.R.A., Moizeau, F., Thisse, J.-F., 2022. Who lives where in the city? Amenities, commuting and income sorting. Journal of Urban Economics 128, 103394. 10.1016/j.jue.2021.103394

Goffette-Nagot, F., Schaeffer, Y., 2013. Accessibilité ou voisinage ?:Une analyse des sources de la ségrégation résidentielle au sein des aires urbaines françaises. Revue économique 64, 857–882. 10.3917/reco.645.0857

González-García, A., Palomo, I., Codemo, A., Rodeghiero, M., Dubo, T., Vallet, A., Lavorel, S., 2025. Co-benefits of nature-based solutions exceed the costs of implementation. Cell Reports Sustainability 2, 100336. 10.1016/j.crsus.2025.100336

González-García, A., Palomo, I., González, J.A., López, C.A., Montes, C., 2020. Quantifying spatial supply-demand mismatches in ecosystem services provides insights for land-use planning. Land Use Policy 94, 104493. 10.1016/j.landusepol.2020.104493

Gronlund, C.J., 2014. Racial and socioeconomic disparities in heat-related health effects and their mechanisms: a review. Curr Epidemiol Rep 1, 165–173. 10.1007/s40471-014-0014-4

Heblich, S., Trew, A., Zylberberg, Y., 2021. East-Side Story: Historical Pollution and Persistent Neighborhood Sorting. Journal of Political Economy 129, 1508–1552. 10.1086/713101

Heckert, M., Rosan, C.D., 2016. Developing a green infrastructure equity index to promote equity planning. Urban Forestry & Urban Greening, Special Section: Power in urban social-ecological systems: Processes and practices of governance and marginalization 19, 263–270. 10.1016/j.ufug.2015.12.011

Herreros-Cantis, P., McPhearson, T., 2021. Mapping supply of and demand for ecosystem services to assess environmental justice in New York City. Ecological Applications 31, e02390. 10.1002/eap.2390

Hijmans, R.J., Bivand, R., Cordano, E., Dyba, K., Pebesma, E., Sumner, M.D., 2024. terra: Spatial Data Analysis.

Ibes, D.C., 2015. A multi-dimensional classification and equity analysis of an urban park system: A novel methodology and case study application. Landscape and Urban Planning 137, 122–137. 10.1016/j.landurbplan.2014.12.014

IGN, 2023. CoSIA [WWW Document]. URL https://cosia.ign.fr/info#descriptif (accessed 12.3.24).

Insee, 2019. Revenus, pauvreté et niveau de vie - Données carroyées. Dispositif Fichier localisé social et fiscal (FiLoSoFi).

Jiang, B., Larsen, L., Deal, B., Sullivan, W.C., 2015. A dose–response curve describing the relationship between tree cover density and landscape preference. Landscape and Urban Planning 139, 16–25. 10.1016/j.landurbplan.2015.02.018

Kabisch, N., Haase, D., 2014. Green justice or just green? Provision of urban green spaces in Berlin, Germany. Landscape and Urban Planning 122, 129–139. 10.1016/j.landurbplan.2013.11.016

Karger, D.N., Conrad, O., Böhner, J., Kawohl, T., Kreft, H., Soria-Auza, R.W., Zimmermann, N.E., Linder, H.P., Kessler, M., 2017. Climatologies at high resolution for the earth’s land surface areas. Sci Data 4, 170122. 10.1038/sdata.2017.122

Kato-Huerta, J., Geneletti, D., 2022. Environmental justice implications of nature-based solutions in urban areas: A systematic review of approaches, indicators, and outcomes. Environmental Science & Policy 138, 122–133. 10.1016/j.envsci.2022.07.034

Kronenberg, J., Haase, A., Łaszkiewicz, E., Antal, A., Baravikova, A., Biernacka, M., Dushkova, D., Filčak, R., Haase, D., Ignatieva, M., Khmara, Y., Niţă, M.R., Onose, D.A., 2020. Environmental justice in the context of urban green space availability, accessibility, and attractiveness in postsocialist cities. Cities 106, 102862. 10.1016/j.cities.2020.102862

Lapointe, M., 2020. Urbanization alters ecosystem service preferences in a Small Island Developing State. Ecosystem Services 9.

Lapointe, M., Gurney, G., Cumming, G., 2020. Perceived availability and access limitations to ecosystem service well-being benefits increase in urban areas. Ecology and Society 25. 10.5751/ES-12012-250432

Law, A., Carrasco, L.R., Richards, D.R., Shaikh, S.F.E.A., Tan, C.L.Y., Nghiem, L.T.P., 2022. Leave no one behind: A case of ecosystem service supply equity in Singapore. Ambio 51, 2118–2136. 10.1007/s13280-022-01735-x

Lejemtel, L., Vallet, A., Chiron, F., Levrel, H., Lavorel, S., 2025. Exploring environmental justice and nexus approaches in Paris urban nature policies. Journal of Environmental Management 392, 126547. 10.1016/j.jenvman.2025.126547

Liotta, C., Kervinio, Y., Levrel, H., Tardieu, L., 2020. Planning for environmental justice - reducing well-being inequalities through urban greening. Environmental Science & Policy 112, 47–60. 10.1016/j.envsci.2020.03.017

Lüdecke, D., 2024. sjstats: Collection of Convenient Functions for Common Statistical Computations.

Margaritis, E., Kang, J., Filipan, K., Botteldooren, D., 2018. The influence of vegetation and surrounding traffic noise parameters on the sound environment of urban parks. Applied Geography 94, 199–212. 10.1016/j.apgeog.2018.02.017

Marselle, M.R., Bowler, D.E., Watzema, J., Eichenberg, D., Kirsten, T., Bonn, A., 2020. Urban street tree biodiversity and antidepressant prescriptions. Sci Rep 10, 22445. 10.1038/s41598-020-79924-5

Marsoner, T., Simion, H., Giombini, V., Egarter Vigl, L., Candiago, S., 2023. A detailed land use/land cover map for the European Alps macro region. Sci Data 10, 468. 10.1038/s41597-023-02344-3

Martinez-Harms, M.J., Bryan, B.A., Wood, S.A., Fisher, D.M., Law, E., Rhodes, J.R., Dobbs, C., Biggs, D., Wilson, K.A., 2018a. Inequality in access to cultural ecosystem services from protected areas in the Chilean biodiversity hotspot. Science of The Total Environment 636, 1128–1138. 10.1016/j.scitotenv.2018.04.353

Martinez-Harms, M.J., Bryan, B.A., Wood, S.A., Fisher, D.M., Law, E., Rhodes, J.R., Dobbs, C., Biggs, D., Wilson, K.A., 2018b. Inequality in access to cultural ecosystem services from protected areas in the Chilean biodiversity hotspot. Science of The Total Environment 636, 1128–1138. 10.1016/j.scitotenv.2018.04.353

Martín-López, B., Iniesta-Arandia, I., García-Llorente, M., Palomo, I., Casado-Arzuaga, I., Amo, D.G.D., Gómez-Baggethun, E., Oteros-Rozas, E., Palacios-Agundez, I., Willaarts, B., González, J.A., Santos-Martín, F., Onaindia, M., López-Santiago, C., Montes, C., 2012. Uncovering Ecosystem Service Bundles through Social Preferences. PLoS ONE 7, e38970. 10.1371/journal.pone.0038970

Meyer-Grandbastien, A., Burel, F., Hellier, E., Bergerot, B., 2020. A step towards understanding the relationship between species diversity and psychological restoration of visitors in urban green spaces using landscape heterogeneity. Landscape and Urban Planning 195, 103728. 10.1016/j.landurbplan.2019.103728

“Natural Capital Project,” 2024. InVEST.

Neier, T., 2023. The green divide: A spatial analysis of segregation-based environmental inequality in Vienna. Ecological Economics 213, 107949. 10.1016/j.ecolecon.2023.107949

Nygaard, C.A., 2024. Green infrastructure and socioeconomic dynamics in London low-income neighbourhoods: A 120-year perspective. Cities 144, 104616. 10.1016/j.cities.2023.104616

“OpenStreetMap contributors,” 2024. OpenStreetMap [Data set]. OpenStreetMap Foundation. Available as open data under the Open Data Commons Open Database License (ODbL) at openstreetmap.org.

Özgüner, H., 2011. Cultural Differences in Attitudes towards Urban Parks and Green Spaces. Landscape Research 36, 599–620.

Pan, H., Page, J., Cong, C., Barthel, S., Kalantari, Z., 2021. How ecosystems services drive urban growth: Integrating nature-based solutions. Anthropocene 35, 100297. 10.1016/j.ancene.2021.100297

Reid, C.E., O’Neill, M.S., Gronlund, C.J., Brines, S.J., Brown, D.G., Diez-Roux, A.V., Schwartz, J., 2009. Mapping community determinants of heat vulnerability. Environ Health Perspect 117, 1730–1736. 10.1289/ehp.0900683

Reyes-Riveros, R., Altamirano, A., De La Barrera, F., Rozas-Vásquez, D., Vieli, L., Meli, P., 2021. Linking public urban green spaces and human well-being: A systematic review. Urban Forestry & Urban Greening 61, 127105. 10.1016/j.ufug.2021.127105

Richards, D., Polyakov, M., Brandt, A.J., Cavanagh, J., Diprose, G., Milner, G., Ramana, J.V., Simcock, R., 2023. Inequity in nature’s contributions to people in ōtautahi/Christchurch: A low-density post-earthquake city. Urban Forestry & Urban Greening 86, 128044. 10.1016/j.ufug.2023.128044

Rigolon, A., 2017. Parks and young people: An environmental justice study of park proximity, acreage, and quality in Denver, Colorado. Landscape and Urban Planning 165, 73–83. 10.1016/j.landurbplan.2017.05.007

Romero, H., Vásquez, A., Fuentes, C., Salgado, M., Schmidt, A., Banzhaf, E., 2012. Assessing urban environmental segregation (UES). The case of Santiago de Chile. Ecological Indicators 23, 76–87. 10.1016/j.ecolind.2012.03.012

Ross, C.W., Prihodko, L., Anchang, J.Y., Kumar, S.S., Ji, W., Hanan, N.P., 2018. Global Hydrologic Soil Groups (HYSOGs250m) for Curve Number-Based Runoff Modeling. 10.3334/ORNLDAAC/1566

Sari, E.N., Bayraktar, S., 2023. The role of park size on ecosystem services in urban environment: a review. Environ Monit Assess 195, 1072. 10.1007/s10661-023-11644-5

Schaeffer, Y., Cremer-Schulte, D., Tartiu, C., Tivadar, M., 2016. Natural amenity-driven segregation: Evidence from location choices in French metropolitan areas. Ecological Economics 130, 37–52. 10.1016/j.ecolecon.2016.05.018

Schaeffer, Y., Tivadar, M., 2019. Measuring Environmental Inequalities: Insights from the Residential Segregation Literature. Ecological Economics 164, 106329. 10.1016/j.ecolecon.2019.05.009

Schlosberg, D., 2013. Theorising environmental justice: the expanding sphere of a discourse. Env. Polit. 22, 37–55. 10.1080/09644016.2013.755387

Tivadar, M., 2019. OasisR: An R Package to Bring Some Order to the World of Segregation Measurement. Journal of Statistical Software 89, 1–39. 10.18637/jss.v089.i07

Tu, G., Abildtrup, J., Garcia, S., 2016. Preferences for urban green spaces and peri-urban forests: An analysis of stated residential choices. Landscape and Urban Planning 148, 120–131. 10.1016/j.landurbplan.2015.12.013

“United Nations,” 2018. World Urbanization Prospects, Population Division - UN.

Vannier, C., Bierry, A., Longaretti, P.-Y., Nettier, B., Cordonnier, T., Chauvin, C., Bertrand, N., Quétier, F., Lasseur, R., Lavorel, S., 2019. Co-constructing future land-use scenarios for the Grenoble region, France. Landscape and Urban Planning 190, 103614. 10.1016/j.landurbplan.2019.103614

Venter, Z.S., Shackleton, C.M., Van Staden, F., Selomane, O., Masterson, V.A., 2020. Green Apartheid: Urban green infrastructure remains unequally distributed across income and race geographies in South Africa. Landscape and Urban Planning 203, 103889. 10.1016/j.landurbplan.2020.103889

Xing, J., Hu, Z., Xia, F., Xu, J., Zou, E., 2023. Urban Forests: Environmental Health Values and Risks. Working Paper Series. 10.3386/w31554

Yasumoto, S., Nakaya, T., Jones, A.P., 2021. Quantitative Environmental Equity Analysis of Perceived Accessibility to Urban Parks in Osaka Prefecture, Japan. Appl. Spatial Analysis 14, 337–354. 10.1007/s12061-020-09360-5

Yi, H., Kreuter, U.P., Han, D., Güneralp, B., 2019. Social segregation of ecosystem services delivery in the San Antonio region, Texas, through 2050. Science of The Total Environment 667, 234–247. 10.1016/j.scitotenv.2019.02.130

Zomer, R.J., Xu, J., Trabucco, A., 2022. Version 3 of the Global Aridity Index and Potential Evapotranspiration Database. Sci Data 9, 409. 10.1038/s41597-022-01493-1

