## Supplementary material for "Disparities in climate risk-mitigation and recreation services in Grenoble (France)"

### Supplementary methods

*Figure S1. Map of the study area*

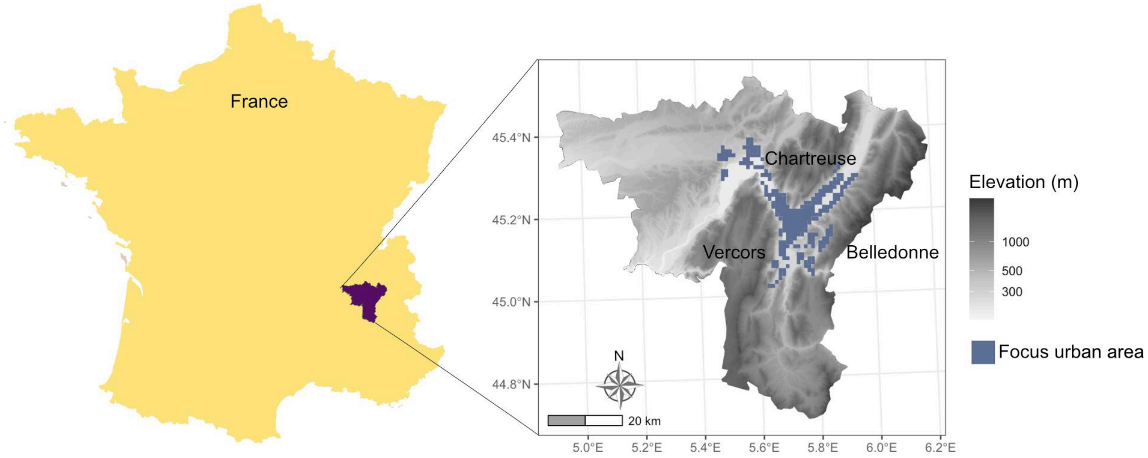


#### Data sources

*Table S1. Description of data sources*

| Data type | Data source | Description |
|---|---|---|
| Land-use land cover | EuSAlps (Marsoner et al., 2023) | Detailed land use-land cover map of the whole alpine region, 5m resolution |
| Land-use land cover | CosIA (IGN, 2023) | Predicted land use land cover map from deep learning approaches, particularly useful for urban land uses. Aggregated to 10m resolution |
| Urban areas delimitation | Global Human Settlement Index (Schiavina et al., 2023) | Raster map describing and classifying settlement typologies, urban or rural, from population density data or build-up areas density, with a 1km resolution.<br>30 : Urban Center grid cell<br>23 : Dense Urban Cluster grid cell<br>22 : Semi-dense Urban Cluster grid cell<br>21 : Suburban or Peri-urban grid cell<br>13 : Low Density Pural grid cell<br>11 : Very Low Density Rural grid cell |
| Green space delimitation, Amenities, roads | (OpenStreetMap contributors, 2024) | Green spaces: areas classified in OpenStreetMap land use as cemeteries, forests, gardens, meadows, nature reserves, parks, village_green and woods<br><br>Amenities: point data classified in OpenStreetMap as leisure\|playgrounds, amenity\|water points, amenity\|drinking_water, and leisure\|fitness stations.<br><br>Roads: Combination of multiple OpenStreetMap layers. Noise level 1: highways classified as tertiary, unclassified, residential, living stree, or service; noise |

| | | |
|---|---|---|
| | | level 2: highways classified as motorways, trunks, or secondary; noise level 3: primary highways and rails |
| Precipitation | Chelsa climate (Karger et al., 2017) | Global precipitation data (resolution 30 arc sec) |
| Soil retention capacity | | |
| Evapotranspiration | Potential Evapotranspiration Database (Zomer et al., 2022) | High-resolution (30 arc-seconds) global hydro-climatic data averaged (1970–2000) monthly and yearly, based upon the FAO Penman-Monteith Reference Evapotranspiration ($ET_0$) equation. |
| Socio-demographic data | Filosofi dataset (Insee, 2019) | Data on the age and economic structure of the population at a 200m resolution |

#### Model parameterisation

Table S2. Land use parameters used for the heat mitigation index model

| EUSALP code | EUSALP description | Albedo | Kc | Green area | Shade | References |
|---|---|---|---|---|---|---|
| **11000** | Artificial surfaces and constructions | 0.143 | 0.325 | 0 | 0 | Trlica et al., 2017 |
| **11100** | Dense settlement area | 0.143 | 0.325 | 0 | 0 | |
| **11200** | Low density settlement area | 0.147 | 0.25 | 0 | 0 | |
| **11300** | Builtup area | 0.143 | 0.325 | 0 | 0 | |
| **11400** | Open settlement area | 0.147 | 0.25 | 0 | 0 | |
| **12100** | Industrial and commercial zones | 0.132 | 0.35 | 0 | 0 | |
| **12210** | Roads motroways and trunks | 0.139 | 0.3 | 0 | 0 | |
| **12220** | Road networks | 0.139 | 0.3 | 0 | 0 | |
| **12221** | Roads tertiary and others | 0.139 | 0.3 | 0 | 0 | |
| **12230** | Railways train tracks | 0.139 | 0.3 | 0 | 0 | |
| **12240** | Unpaved roads and tracks | 0.132 | 0.31 | 0 | 0 | |
| **14100** | Green urban areas | 0.136 | 0.27 | 1 | 0 | average (311, 312, 313, 320, 321) |
| **21000** | Cultivated areas - Arable land - Annual crops | 0.180 | 0.912 | 1 | 0 | Average annual crops |
| **21211** | Common wheat | 0.186 | 0.89 | 1 | 0 | Kang et al., 2003; Sieber et al., 2022 |
| **21212** | Durum wheat | 0.186 | 0.8 | 1 | 0 | Lhomme et al., 2009 |

| | | | | | | |
|---|---|---|---|---|---|---|
| **21213** | Barley | 0.176 | 1.07 | 1 | 0 | Attarod et al., 2009; Sieber et al., 2022 |
| **21214** | Rye | 0.181 | 0.87 | 1 | 0 | Sieber et al. 2022 |
| **21215** | Oats | 0.172 | 0.867 | 1 | 0 | Andréasson, 2023; Moteva et al., 2014 |
| **21216** | Maize | 0.184 | 0.973 | 1 | 0 | Bsaibes et al., 2009; Kang et al., 2003 |
| **21218** | Triticale | 0.183 | 0.978 | 1 | 0 | Moteva et al. 2014 |
| **21221** | Potatoes | 0.214 | 0.65 | 1 | 0 | Paredes et al., 2018 |
| **21222** | Sugar beet | 0.179 | 0.918 | 1 | 0 | Sieber et al. 2022, Moteva et al. 2014 |
| **21230** | Other non-permanent industrial crops | 0.180 | 0.912 | 1 | 0 | Same as 21000 |
| **21231** | Sunflower | 0.225 | 0.79 | 1 | 0 | Mila et al., 2016; Srivastava et al., 1998 |
| **21232** | Rape and turnip rape | 0.188 | 0.945 | 1 | 0 | Sieber et al. 2022, Moteva et al. 2014 |
| **21233** | Soya | 0.239 | 0.86 | 1 | 0 | Weiss et al. 2001, Attarod et al. 2009 |
| **21240** | Dry pulses | 0.214 | 0.8 | 1 | 0 | Nandi et al., 2024 |
| **21250** | Fodder crops (cereals and leguminous) | 0.214 | 0.884 | 1 | 0 | Weiss et al. 2001, Moteva et al. 2014 |
| **21290** | Bare arable land | 0.23 | 0.275 | 1 | 0 | Same as 33100 |
| **22000** | Permanent crops | 0.115 | 1.075 | 1 | 0.7 | Schwaab et al., 2015 |
| **22100** | Vineyard | 0.2 | 0.86 | 1 | 0 | Cancela et al., 2010; Galleguillos et al., 2011 |
| **22200** | Orchard | 0.179 | 0.918 | 1 | 0 | Sieber et al. 2022, Moteva et al. 2014 |
| **23100** | Managed grassland - Pastures | 0.196 | 0.93 | 1 | 0 | Ambrosi et al., 2024; Rosset et al., 2001 |
| **23200** | Seminatural grassland - Meadows | 0.168 | 0.93 | 1 | 0 | |
| **31100** | Broadleaf tree cover | 0.126 | 1.55 | 1 | 1 | |
| **31102** | Broadleaf tree cover 30-60% | 0.126 | 1.55 | 1 | 0.45 | |
| **31103** | Broadleaf tree cover 60-100% | 0.126 | 1.55 | 1 | 0.8 | |

| | | | | | | |
|---|---|---|---|---|---|---|
| **31200** | Coniferous tree cover | 0.104 | 1 | 1 | 1 | Schwaab et al., 2015 |
| **31202** | Coniferous tree cover 30-60% | 0.104 | 1 | 1 | 0.45 | |
| **31203** | Coniferous tree cover 60-100% | 0.104 | 1 | 1 | 0.8 | |
| **31300** | Mixed tree cover | 0.115 | 1.4 | 1 | 1 | |
| **31400** | Tree cover in agricultural context | 0.115 | 1.075 | 1 | 0.7 | |
| **31450** | Tree cover in urban context | 0.136 | 0.27 | 1 | 0.7 | Same as 14100 |
| **31500** | Green linear elements - linear woody features | 0.132 | 0.975 | 1 | 0.4 | Aartsma et al., 2020 |
| **31600** | Patchy woody features | 0.132 | 0.975 | 1 | 0 | |
| **31610** | Additional woody features | 0.132 | 0.975 | 1 | 0 | |
| **32000** | Scrub and shrubland | 0.132 | 0.975 | 1 | 0 | |
| **32100** | Alpine and sub-alpine natural grassland | 0.194 | 1.125 | 1 | 0 | Blumthaler and Ambach, 1988; Tian et al., 2014 |
| **32200** | Moors and heathland - other scrubland | 0.132 | 0.975 | 1 | 0 | Aartsma et al., 2020 |
| **32300** | Sclerophyllous vegetation | 0.132 | 0.65 | 1 | 0 | |
| **33100** | Beaches, dunes, sands | 0.23 | 0.275 | 1 | 0 | Blumthaler and Ambach, 1988 |
| **33200** | Bare rocks and rock debris | 0.3 | 0.125 | 1 | 0 | Nasif Al Fahdawi et al., 2021 |
| **33300** | Sparsely vegetated land | 0.13 | 0.55 | 1 | 0 | Sieber et al., 2022 |
| **33500** | Permanent snow covered surfaces | 0.507 | 0.95 | 1 | 0 | Kalitin, 1930 |
| **41000** | Wetland (permanent wet areas) - inland marshes | 0.159 | 0.625 | 1 | 0 | Eichelmann et al., 2018; Trlica et al., 2017 |
| **51000** | Water bodies | 0.091 | 0.95 | 1 | 0 | Blumthaler and Ambach, 1988 |
| **51100** | Rivernetwork | 0.091 | 0.95 | 1 | 0 | |
| **51200** | Riverbed > 10m width | 0.091 | 0.95 | 1 | 0 | |

Table S3. Land use parameters used in the flood mitigation model

| **EUSALP code** | **EUSALP description** | **cn_a** | **cn_b** | **cn_c** | **cn_d** | **Reference** | **Comment** |
|---|---|---|---|---|---|---|---|
| **11000** | Artificial surfaces and constructions | 98 | 98 | 98 | 98 | Cronshey, 1986 | Impervious areas |
| **11100** | Dense settlement area | 98 | 98 | 98 | 98 | | |
| **11200** | Low density settlement area | 61 | 75 | 83 | 87 | | ¼ acre (38% imp.) |
| **11300** | Builtup area | 98 | 98 | 98 | 98 | | Impervious areas |
| **11400** | Open settlement area | 61 | 75 | 83 | 87 | | ¼ acre (38% imp.) |
| **12100** | Industrial and commercial zones | 98 | 98 | 98 | 98 | | Impervious areas |

| | | | | | | | |
|---|---|---|---|---|---|---|---|
| **12210** | Roads motroways and trunks | 98 | 98 | 98 | 98 | | |
| **12220** | Road networks | 98 | 98 | 98 | 98 | | |
| **12221** | Roads tertiary and others | 98 | 98 | 98 | 98 | | |
| **12230** | Railways train tracks | 98 | 98 | 98 | 98 | | |
| **12240** | Unpaved roads and tracks | 77 | 86 | 91 | 94 | | Bare soil |
| **14100** | Green urban areas | 39 | 61 | 74 | 80 | | Open space (Good condition) |
| **21000** | Cultivated areas - Arable land - Annual crops | 60 | 72 | 80 | 84 | | Small grain SR+CR good |
| **21211** | Common wheat | 60 | 72 | 80 | 84 | | |
| **21212** | Durum wheat | 60 | 72 | 80 | 84 | | |
| **21213** | Barley | 60 | 72 | 80 | 84 | | |
| **21214** | Rye | 60 | 72 | 80 | 84 | | |
| **21215** | Oats | 60 | 72 | 80 | 84 | | |
| **21216** | Maize | 65 | 75 | 82 | 86 | | Row crops contoured |
| **21218** | Triticale | 60 | 72 | 80 | 84 | | Small grain SR+CR good |
| **21221** | Potatoes | 32 | 58 | 72 | 79 | | Orchard |
| **21222** | Sugar beet | 32 | 58 | 72 | 79 | | |
| **21230** | Other non permanent industrial crops | 60 | 72 | 80 | 84 | | Small grain SR+CR good |
| **21231** | Sunflower | 64 | 75 | 82 | 85 | Cronshey 1986 | Row crops SR+CR |
| **21232** | Rape and turnip rape | 64 | 75 | 82 | 85 | | Close-seeded or broadcast legumes or rotation meadow |
| **21233** | Soya | 58 | 72 | 81 | 85 | | |
| **21240** | Dry pulses | 58 | 72 | 81 | 85 | | |
| **21250** | Fodder crops (cereals and leguminous) | 58 | 72 | 81 | 85 | | |
| **21290** | Bare arable land | 77 | 86 | 91 | 94 | | Bare arable land |
| **22000** | Permanent crops | 43 | 65 | 76 | 82 | | Agro-forestry areas |
| **22100** | Vinyard | 67 | 78 | 85 | 89 | | Straight row |
| **22200** | Orchard | 32 | 58 | 72 | 79 | | Orchard |
| **23100** | Managed grassland - Pastures | 8 | 79 | 86 | 89 | | Pasture |
| **23200** | Seminatural grassland - Meadows | 30 | 58 | 71 | 78 | | Meadow |
| **31100** | Broadleaf tree cover | 30 | 55 | 70 | 77 | | Good condition |
| **31102** | Broadleaf tree cover 30-60% | 36 | 60 | 73 | 79 | | Fair condition |

| | | | | | | | |
|---|---|---|---|---|---|---|---|
| **31103** | Broadleaf tree cover 60-100% | 30 | 55 | 70 | 77 | Jaafar et al., 2019; Tedela et al., 2012 | Good condition |
| **31200** | Coniferous tree cover | 33 | 58 | 72 | 78 | | Good condition |
| **31202** | Coniferous tree cover 30-60% | 40 | 63 | 75 | 80 | | Fair condition |
| **31203** | Coniferous tree cover 60-100% | 33 | 58 | 72 | 78 | | Good condition |
| **31300** | Mixed tree cover | 31.5 | 56.5 | 71 | 77.5 | | Average broadleaf and conifer |
| **31400** | Tree cover in agricultural context | 43 | 65 | 76 | 82 | Cronshey 1986 | Woods—grass combination (orchard or tree farm).D (Fair) |
| **31450** | Tree cover in urban context | 39 | 61 | 74 | 80 | | Open space (Good condition) |
| **31500** | Green linear elements - linear woody features | 43 | 65 | 76 | 82 | | Woods—grass combination (orchard or tree farm).D |
| **31600** | Patchy woody features | 43 | 65 | 76 | 82 | | |
| **31610** | Additional woody features | 43 | 65 | 76 | 82 | | |
| **32000** | Scrub and shrubland | 43 | 65 | 76 | 82 | | |
| **32100** | Alpine and sub-alpine natural grassland | 39 | 61 | 74 | 80 | | Brush—brush-weed-grass mixture with brush the major element.B (Fair) |
| **32200** | Moors and heathland - other scrubland | 35 | 56 | 70 | 77 | | Brush—brush-weed-grass mixture with brush the major element.B |
| **32300** | Sclerophyllous vegetation | 0 | 62 | 74 | 85 | | Herbaceous for arid rangelands |
| **33100** | Beaches, dunes, sands | 63 | 77 | 85 | 88 | | Natural desert landscaping (pervious areas only) |
| **33200** | Bare rocks and rock debris | 63 | 77 | 85 | 88 | | |
| **33300** | Sparsely vegetated land | 68 | 79 | 86 | 89 | | Poor condition (grass cover < 50%) |
| **33500** | Permanent snow covered surfaces | 99 | 99 | 99 | 99 | | |
| **41000** | Wetland (permanent wet areas) - inland marshes | 49 | 69 | 79 | 84 | Chen et al., 2014 | Wet tussock grassland with herbs, sedges or rushes, herblands or ferns |
| **51000** | Water bodies | 99 | 99 | 99 | 99 | | |
| **51100** | Rivernetwork | 99 | 99 | 99 | 99 | | |
| **51200** | Riverbed > 10m width | 99 | 99 | 99 | 99 | | |

#### Gini-based environmental segregation indices

The segregation Gini coefficient is a spatial adaptation of the Gini inequality index (Gini, 1921) by Duncan and Duncan (1955). The index compares the distribution of a group relative to another group by adapting the Lorenz inequality curve (Lorenz, 1905) to a spatial context. To construct the segregation curve, spatial units are ranked from lowest to highest according to the share of the first group within each spatial unit and the cumulative proportions of both groups are calculated. Graphically Gini index represents the area between the even diagonal and the segregation curve divided by the area below the diagonal. The coefficient ranges from 0 (complete evenness, identical relative spatial distribution) to 1 (complete separation).

Here we apply the Gini coefficient to environmental segregation: the **Gini Based Environmental Scores Inequality Index**. It measures the degree of unevenness of the relative spatial distribution of two groups, compared to the distribution of an amenity score *a*. The spatial units are ordered following a decrease in the amenity score and the index ranges between -1 and 1: negative values indicate that the first group (x) is more segregated to the amenity than the second group (y) and conversely.

$$GINIes^{x,y} = \left(\sum_{i=2}^{n} X_{i-1}Y_{i}\right) - \left(\sum_{i=2}^{n} X_{i}Y_{i-1}\right)$$

where, *n* is the number of spatial units and $X_i$ and $Y_i$ are the cumulative percentage of each group population through the *i*th spatial unit, spatial units being ordered decreasing by the amenity score *a*.

GINIes follow desirable properties:

- It is negative when weighted mean of group is lower than the mean of y and vice-versa
- Its absolute value is proportional to the absolute value of the means difference (high difference in means = high GINIes)
- Its absolute value it is bounded by GINI segregation index
- Its signs change when compute the index for 1-*a* (the inversed environmental score)

### Supplementary results

***Figure S2.*** *Map of the proportion of vegetation in a 300m radius.*

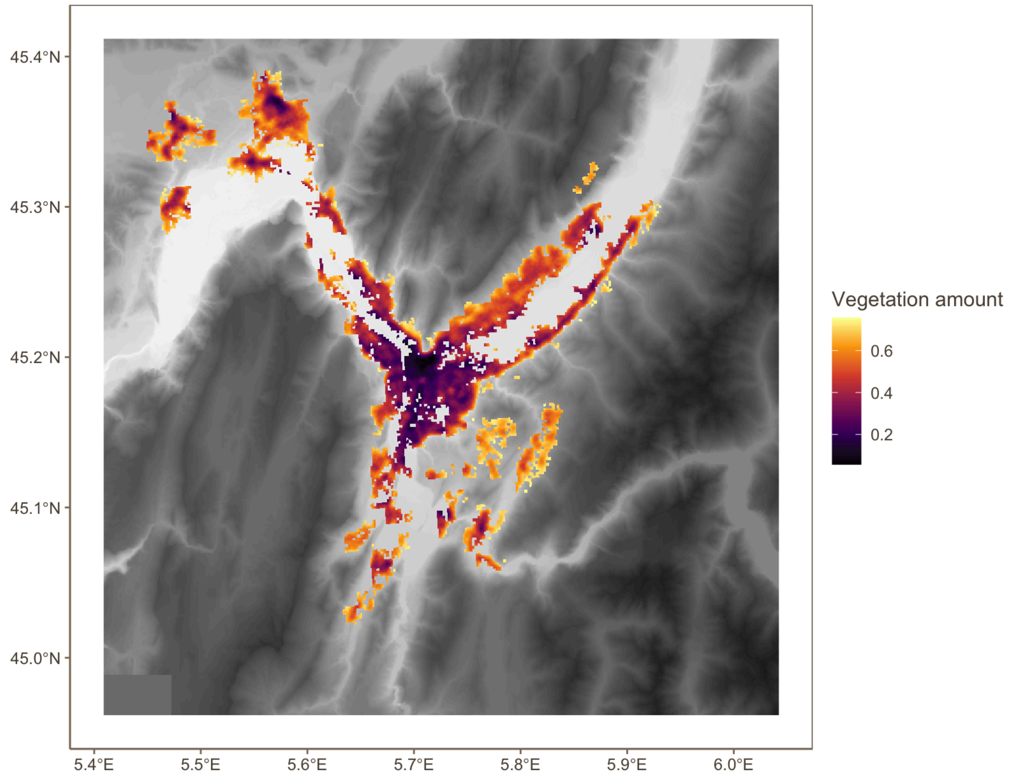


#### Spatial regressions

We complemented our main analyses by spatial regressions, to account for the fact that lower services are provided especially in areas with higher population density and lower economic status. Consistently with the main results, heat and flood mitigation decreased with the proportion of poor households ($\beta_{hmi,\ low}$ = -0.017 ± 0.003 and $\beta_{retention,\ low}$ = -0.14 ± 0.02, $P < 0.001$) and increased with average living standard ($\beta_{hmi,\ living\ std}$ = 0.006 ± 0.001 and $\beta_{retention,\ living\ std}$ = 0.045 ± 0.004, $P < 0.001$). Urban recreation did not vary with the proportion of poor households ($P = 0.41$) but slightly decreased with living standard ($\beta_{park\ quality,\ living\ std}$ = -0.006 ± 0.002, $P < 0.001$) while the distance to the closest park was slightly lower in areas with lower more poor households and lower living standards ($\beta_{distance,\ low}$ = -36.9 ± 9.1 and $\beta_{distance,\ living\ standard}$ = 8.6 ± 2.1, $P < 0.001$) (Table S2).

***Table S2.*** *Spatial regressions with ecosystem service or environmental amenity provided with the proportion of poor households ang average living standard per individual. Regressions were run on all available 200m squares (n = 4801) weighted by total population density.*

| **Ecosystem service** | **ES variation with proportion of poor households (spatial regression with population density-based weights)** | **ES variation with average standard of living (spatial regression with population density-based weights)** |
|---|---|---|
| **HMI** | Slope estimate: -0.017 ± 0.003<br>P-value < 0.001 | Slope estimate: 0.006 ± 0.001<br>P-value < 0.001 |
| **Flood mitigation** | Slope estimate: -0.14 ± 0.02<br>P-value < 0.001 | Slope estimate: 0.045 ± 0.004<br>P-value < 0.001 |
| **Quality of best greenspace within 500m** | Slope estimate: 0.006 ± 0.007<br>P-value = 0.41 | Slope estimate: -0.006 ± 0.002<br>P-value < 0.001 |
| **Distance** | Slope estimate: -36.9 ± 9.1<br>P-value < 0.001 | Slope estimate: 8.6 ± 2.1<br>P-value < 0.001 |

### Supplementary references


Aartsma, P., Asplund, J., Odland, A., Reinhardt, S., Renssen, H., 2020. Surface albedo of alpine lichen heaths and shrub vegetation. Arctic, Antarctic, and Alpine Research 52, 312–322. https://doi.org/10.1080/15230430.2020.1778890

Ambrosi, L., Berger, V., Rainer, G., Obojes, N., Tappeiner, U., Tasser, E., Leitinger, G., 2024. Spatiotemporal variability of evapotranspiration in Alpine grasslands and its biotic and abiotic drivers. Ecohydrology 17, e2633. https://doi.org/10.1002/eco.2633

Andréasson, V., 2023. Evaluating the impact of agriculture on albedo using Sentinel-2 data in southern Sweden. Student thesis series INES.

Attarod, P., Aoki, M., Bayramzadeh, V., 2009. Measurements of the actual evapotranspiration and crop coefficients of summer and winter seasons crops in Japan. Plant, Soil and Environment 55, 121–127. https://doi.org/10.17221/324-PSE

Blumthaler, M., Ambach, W., 1988. Solar Uvb-Albedo of Various Surfaces. Photochemistry and Photobiology 48, 85–88. https://doi.org/10.1111/j.1751-1097.1988.tb02790.x

Bsaibes, A., Courault, D., Baret, F., Weiss, M., Olioso, A., Jacob, F., Hagolle, O., Marloie, O., Bertrand, N., Desfond, V., Kzemipour, F., 2009. Albedo and LAI estimates from FORMOSAT-2 data for crop monitoring. Remote Sensing of Environment 113, 716–729. https://doi.org/10.1016/j.rse.2008.11.014

Cancela, J.J., Fandino, M., Rey, B.J., Pereira, L.S., 2010. Calibration of vineyard crop coefficients and yield response factor to support precision irrigated viticulture. Presented at the International Conference on Agricultural Engineering - AgEng 2010: towards environmental technologies, Clermond-Ferrand, France.

Chen, Y., Wang, B., Pollino, C.A., Cuddy, S.M., Merrin, L.E., Huang, C., 2014. Estimate of flood inundation and retention on wetlands using remote sensing and GIS. Ecohydrology 7, 1412–1420. https://doi.org/10.1002/eco.1467

Cronshey, R., 1986. Urban hydrology for small watersheds (No. 55). US Department of Agriculture, Soil Conservation Service, Engineering Division.

Duncan, O.D., Duncan, B., 1955. Residential Distribution and Occupational Stratification. American Journal of Sociology 60, 493–503.

Eichelmann, E., Hemes, K.S., Knox, S.H., Oikawa, P.Y., Chamberlain, S.D., Sturtevant, C., Verfaillie, J., Baldocchi, D.D., 2018. The effect of land cover type and structure on evapotranspiration from agricultural and wetland sites in the Sacramento–San Joaquin River Delta, California. Agricultural and Forest Meteorology 256–257, 179–195. https://doi.org/10.1016/j.agrformet.2018.03.007

Galleguillos, M., Jacob, F., Prévot, L., French, A., Lagacherie, P., 2011. Comparison of two temperature differencing methods to estimate daily evapotranspiration over a Mediterranean vineyard watershed from ASTER data. Remote Sensing of Environment 115, 1326–1340. https://doi.org/10.1016/j.rse.2011.01.013

Gini, C., 1921. Measurement of Inequality of Incomes. The Economic Journal 31, 124–126. https://doi.org/10.2307/2223319

IGN, 2023. CoSIA [WWW Document]. URL https://cosia.ign.fr/info#descriptif (accessed 12.3.24).

Insee, 2019. Revenus, pauvreté et niveau de vie - Données carroyées. Dispositif Fichier localisé social et fiscal (FiLoSoFi).

Jaafar, H.H., Ahmad, F.A., El Beyrouthy, N., 2019. GCN250, new global gridded curve numbers for hydrologic modeling and design. Sci Data 6, 145. https://doi.org/10.1038/s41597-019-0155-x

Kalitin, N.N., 1930. THE MEASUREMENTS OF THE ALBEDO OF A SNOW COVER. Monthly Weather Review 58, 59–61. https://doi.org/10.1175/1520-0493(1930)58<59:TMOTAO>2.0.CO;2

Kang, S., Gu, B., Du, T., Zhang, J., 2003. Crop coefficient and ratio of transpiration to evapotranspiration of winter wheat and maize in a semi-humid region. Agricultural

Water Management 59, 239–254. https://doi.org/10.1016/S0378-3774(02)00150-6
Karger, D.N., Conrad, O., Böhner, J., Kawohl, T., Kreft, H., Soria-Auza, R.W., Zimmermann, N.E., Linder, H.P., Kessler, M., 2017. Climatologies at high resolution for the earth's land surface areas. Sci Data 4, 170122. https://doi.org/10.1038/sdata.2017.122
Lhomme, J.P., Mougou, R., Mansour, M., 2009. Potential impact of climate change on durum wheat cropping in Tunisia. Climatic Change 96, 549–564. https://doi.org/10.1007/s10584-009-9571-9
Marsoner, T., Simion, H., Giombini, V., Egarter Vigl, L., Candiago, S., 2023. A detailed land use/land cover map for the European Alps macro region. Sci Data 10, 468. https://doi.org/10.1038/s41597-023-02344-3
Mila, A.J., Akanda, A.R., Biswas, S.K., Ali, M.H., 2016. Crop Co-efficient Values of Sunflower for Different Growth Stages by Lysimeter Study. International Journal of Environment and Climate Change 53–63. https://doi.org/10.9734/BJECC/2016/24246
Moteva, M., Kazandjiev, V., Zhivkov, Z., Kireva, R., Mladenova, B., Matev, A., Kalaydzhieva, R., 2014. Estimation of crop evapotranspiration in Bulgarian climate conditions.
Nandi, R., Mudi, D.K., Singh, Kh.C., Saha, M., Bandyopadhyay, P.K., 2024. Partitioning of Evapotranspiration and Crop Coefficients of Lentil Under Conserved Soil Moisture Conditions. J Soil Sci Plant Nutr 24, 435–450. https://doi.org/10.1007/s42729-023-01554-3
Nasif Al Fahdawi, Y.M., Mashee Al Ramahi, F.K., Hamadi Alfalahi, A.S., 2021. Measurement Albedo Coefficient For Land Cover (Lc) And Land Use (Lu), Using Remote Sensing Techniques, A Study Case: Fallujah City. J. Phys.: Conf. Ser. 1829, 012003. https://doi.org/10.1088/1742-6596/1829/1/012003
"OpenStreetMap contributors," 2024. OpenStreetMap [Data set]. OpenStreetMap Foundation. Available as open data under the Open Data Commons Open Database License (ODbL) at openstreetmap.org.
Paredes, P., D'Agostino, D., Assif, M., Todorovic, M., Pereira, L.S., 2018. Assessing potato transpiration, yield and water productivity under various water regimes and planting dates using the FAO dual Kc approach. Agricultural Water Management 195, 11–24. https://doi.org/10.1016/j.agwat.2017.09.011
Rosset, M., Montani, M., Tanner, M., Fuhrer, J., 2001. Effects of abandonment on the energy balance and evapotranspiration of wet subalpine grassland. Agriculture, Ecosystems & Environment 86, 277–286. https://doi.org/10.1016/S0167-8809(00)00290-5
Schiavina, M., Melchiorri, M., Freire, S., 2023. GHS-DUC R2023A - GHS Degree of Urbanisation Classification, application of the Degree of Urbanisation methodology (stage II) to GADM 4.1 layer, multitemporal (1975-2030). European Commission, Joint Research Centre (JRC). https://10.2905/DC0EB21D-472C-4F5A-8846-823C50836305
Schwaab, J., Bavay, M., Davin, E., Hagedorn, F., Hüsler, F., Lehning, M., Schneebeli, M., Thürig, E., Bebi, P., 2015. Carbon storage versus albedo change: radiative forcing of forest expansion in temperate mountainous regions of Switzerland. Biogeosciences 12, 467–487. https://doi.org/10.5194/bg-12-467-2015
Sieber, P., Ericsson, N., Hammar, T., Hansson, P.-A., 2022. Albedo impacts of current agricultural land use: Crop-specific albedo from MODIS data and inclusion in LCA of crop production. Science of The Total Environment 835, 155455. https://doi.org/10.1016/j.scitotenv.2022.155455
Srivastava, N.U., Das, N., Victor, U., Vijaya Kumar, P., Vittal, K.P.R., Ramana Rao, B.V., 1998. Influence of weather parameters on radiation and water efficiency of sunflower (Helianthus annus L.). Indian Journal of Dryland Agriculture Research and Development 12, 55–63.
Tedela, N.H., McCutcheon, S.C., Rasmussen, T.C., Hawkins, R.H., Swank, W.T., Campbell, J.L., Adams, M.B., Jackson, C.R., Tollner, E.W., 2012. Runoff curve numbers for 10 small forested watersheds in the mountains of the eastern United States. Journal of Hydrologic Engineering. 17: 1188-1198. 17, 1188–1198. https://doi.org/10.1061/(ASCE)HE.1943-5584.0000436

Tian, L., Zhang, Y., Zhu, J., 2014. Decreased surface albedo driven by denser vegetation on the Tibetan Plateau. Environ. Res. Lett. 9, 104001. https://doi.org/10.1088/1748-9326/9/10/104001

Trlica, A., Hutyra, L.R., Schaaf, C.L., Erb, A., Wang, J.A., 2017. Albedo, Land Cover, and Daytime Surface Temperature Variation Across an Urbanized Landscape. Earth's Future 5, 1084–1101. https://doi.org/10.1002/2017EF000569

Zomer, R.J., Xu, J., Trabucco, A., 2022. Version 3 of the Global Aridity Index and Potential Evapotranspiration Database. Sci Data 9, 409. https://doi.org/10.1038/s41597-022-01493-1
